# Repression of lysosomal transcription factors Tfeb and Tfe3 is essential for the migration and function of microglia

**DOI:** 10.1101/2020.10.27.357897

**Authors:** Harini Iyer, Kimberle Shen, Ana M. Meireles, William S. Talbot

**Affiliations:** Department of Developmental Biology, Stanford University School of Medicine, Stanford CA 94305

**Author notes:** Genentech Inc, 1 DNA Way, South San Francisco, CA 94080.

## Abstract

As the primary phagocytic cells of the central nervous system, microglia exquisitely regulate their lysosomal activity to facilitate brain development and homeostasis. However, mechanisms that coordinate lysosomal activity with microglia development, migration, and function remain unclear. Here we show that embryonic macrophages require the lysosomal GTPase RagA and the GTPase-activating protein Folliculin to colonize the brain in zebrafish. We demonstrate that embryonic macrophages in *rraga* mutants show increased expression of lysosomal genes but display significant downregulation of immune and migration-related genes. Furthermore, we find that RagA and Folliculin repress the key lysosomal transcription factor Tfeb, and its homologs Tfe3a and Tfe3b, in the macrophage lineage. Using RNA-Sequencing, we establish that Tfeb and Tfe3 are required for activation of lysosomal target genes under conditions of stress but not for basal expression of lysosomal pathways. Collectively, our data define a lysosomal regulatory circuit essential for macrophage development and function in vivo.

**Teaser:** The degradation machinery of the cell must be carefully controlled for the normal formation and function of key immune cells called microglia.

## Introduction

Microglia, the primary resident macrophages of the central nervous system, govern multiple aspects of brain architecture and function. During development, microglia promote synapse formation and maturation, stimulate neurogenesis, and eliminate excess or apoptotic cells (*1-4*). To maintain adult brain homeostasis, microglia surveil the brain for aberrations, mediate repair and regeneration, and facilitate immune clearance of pathogens and cellular debris (*5-10*). As professional phagocytic cells, nearly all these activities of microglia require lysosomes for either degradation of engulfed material or mounting immune responses. Moreover, many studies have found that aberrant microglial activity is associated with neurodegeneration and lysosomal dysregulation (*11-13*). Much less is known about the mechanisms that regulate lysosomes during microglia development. A better understanding of genes regulating lysosomal function during development and homeostasis has the potential to illuminate how these critical organelles might be disrupted during aging and disease.

Central to lysosomal functions are Transcription factor EB (Tfeb) and other members of the Microphthalmia-associated transcription factor (MiTF) protein family. In cell culture and some cell types in vivo, Tfeb, and the related transcription factor Tfe3, have been reported to activate diverse lysosomal processes, such as lysosomal biogenesis, autophagy, and exocytosis (*14-18*). In recent years, there has been an increasing appreciation of the dysregulation of Tfeb in neurodegenerative diseases (*19-23*), which has led to attempts towards enhancing cellular clearance by overexpressing Tfeb to activate autophagy and lysosomal pathways in cell culture or animal models of neurodegeneration (*22, 24-29*). Although exogenous expression of Tfeb improves behavioral deficits and reduces pathology due to misfolded proteins in some models of neurodegenerative diseases (*27-29*), the identity of cells that must overexpress Tfeb to generate these beneficial effects is unclear. Furthermore, the role of Tfeb in microglia, cells that depend on lysosomal pathways for executing many of their functions, is unknown.

The importance of investigating the development and function of microglia in vivo is evident from experiments showing that microglia rapidly lose their transcriptional and epigenetic identity when separated from their niche (*30-34*). Accordingly, studies of microglia in zebrafish using live imaging and mutational studies have led to many important insights into the development and function of these critical glial cells (*35-40*). Here we capitalize on the experimental advantages of zebrafish to functionally define a lysosomal regulatory circuit that is essential in development and function of microglia. We show that the lysosomal GTPase RagA (encoded by *rraga*) is necessary for the normal development, migration, and function of both early microglia and peripheral macrophages. We demonstrate that macrophages in *rraga* mutants significantly upregulate lysosomal genes, but downregulate immune and migration-related genes, revealing a previously unknown role for RagA in the innate immune system. We provide genetic evidence that the lysosomal GTPase activating protein Folliculin is functionally coupled to RagA in macrophages and microglia. Furthermore, we show that simultaneous loss of *tfeb* and *tfe3* (including both zebrafish paralogs of *tfe3, tfe3a* and *tfe3b*) is sufficient to restore microglia numbers in *rraga* and *flcn* mutants; correspondingly, overexpression of *tfe3b* in the macrophage lineage recapitulates all the *rraga* or *flcn* microglia phenotypes we examined. Finally, we uncover that Tfeb and Tfe3 are dispensable for basal levels of lysosomal gene expression, but demonstrate that these transcription factors are required to activate lysosomal pathways in response to stress. Together, our observations reveal a lysosomal pathway that regulates development and function of microglia and macrophages in vivo. Additionally, our observation that Tfeb and Tfe3 must be repressed for normal development and function of microglia has important implications for therapeutic strategies that modulate Tfeb activity in the context of neurodegeneration.

## Results

### Embryonic macrophages fail to colonize the brain in *rraga* mutants

We have previously shown that zebrafish mutants for the gene *rraga*, which encodes the lysosomal GTPase RagA, have fewer microglia than their wildtype siblings (*41*). Previous studies have not, however, addressed the developmental timing or cause of this reduction in microglial cell number in *rraga* mutants. To define the disrupted developmental processes underlying the *rraga* mutant phenotype, we first examined the specification of embryonic macrophages that give rise to microglia. We quantified these cells in embryos between 24 and 26 hours post fertlization (hpf), using the *Tg(mpeg:GFP)* transgene, which expresses green fluorescent protein in the macrophage lineage (*42*). We observed no significant difference in the number of GFP-labeled cells between *rraga* mutants and their wildtype siblings either in the yolk, where these macrophages are initially specified (*43, 44*) (Fig. 1A, 1B), or the rest of the body (Fig. 1A, 1C), indicating that specification and migration of embryonic macrophages is normal at this stage in *rraga* mutants.

**FIG. 1:**
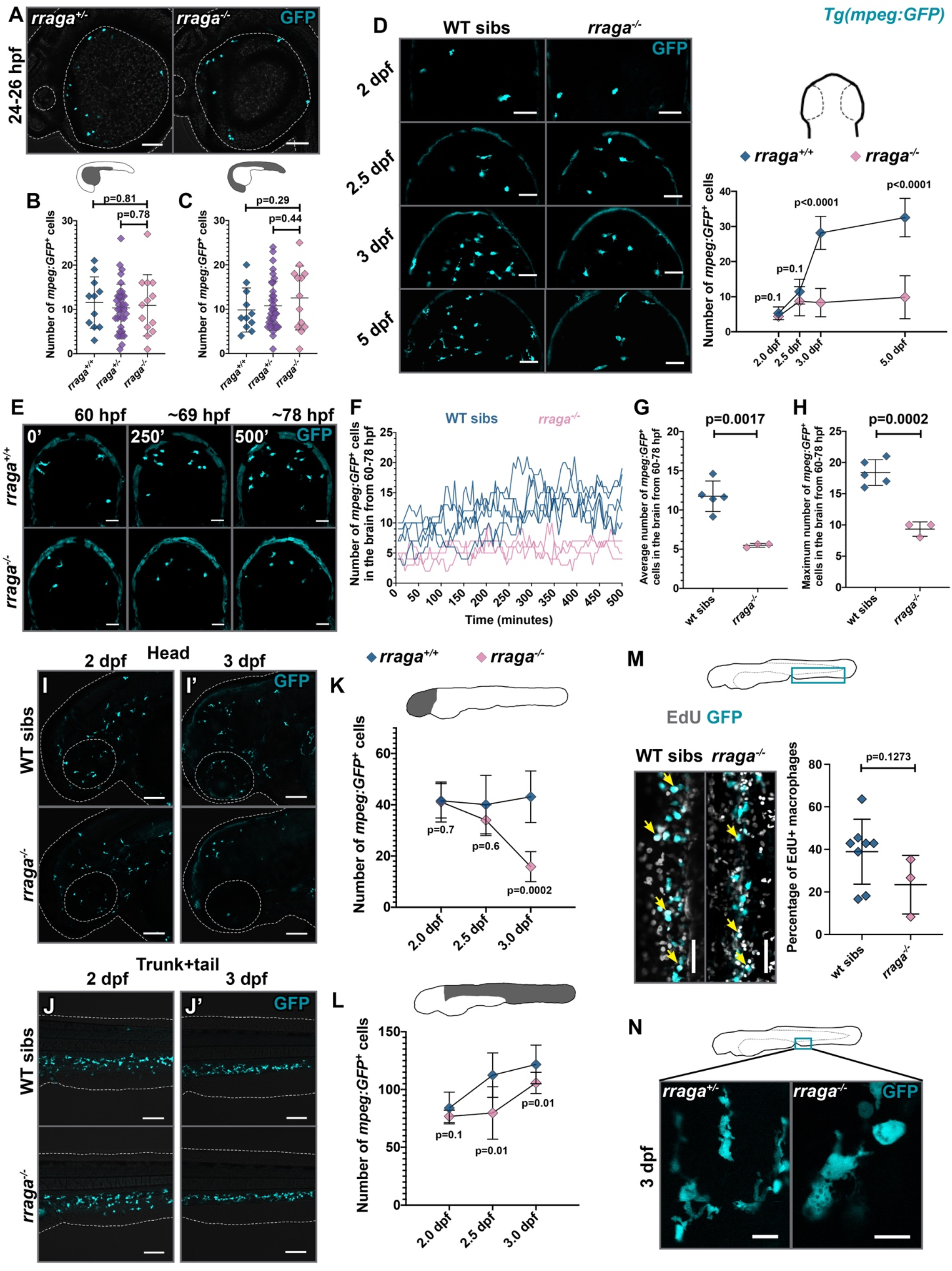
Defective macrophages in *rraga* mutants fail to colonize the brain. (A) images of *mpeg:GFP* expression in *rraga* mutant and heterozygous embryos between 24-26 hpf (lateral views of the head and yolk). Quantitation of macrophages in (B) yolk and (C) rest of the body. (D) Dorsal views of the midbrain showing *mpeg:GFP* expression at the indicated stages along with the corresponding quantification of developing microglia in the brain. (E-H) Timelapse imaging of *mpeg:GFP*^*+*^ macrophages in the midbrain in *rraga* mutants and their siblings between 60 and ∼78 hpf. (E) Representative images from timelapse movies (dorsal views, anterior on top). Images show maximum intensity projections of z-slices at indicated times. (F-H) Quantitative analysis of timelapse movies. In (F), each line shows macrophage cell counts from a single embryo over time. Lateral views of peripheral macrophages and quantitation, (I, I’, K) in the head and (J, J’, L) trunk+tail in *rraga* mutants and their siblings. (M) EdU labeling to determine the percentage of proliferating macrophages in *rraga* mutants and wildtype siblings. Arrows show colocalization of EdU label with macrophages. (N) High magnification images showing the difference in the morphology of macrophages between *rraga* mutant and heterozygous animals at 3 dpf. Scale bars, 50 µm. All graphs show mean with standard deviation; significance was determined using unpaired t-test.

To define the developmental window during when reduction of microglia numbers becomes apparent in *rraga* mutants, we quantified macrophages in the dorsal brain, where microglia are easily visualized, at 2, 2.5, 3, and 5 days post fertilization (dpf) (Fig. 1D). There was a significant reduction in the number of developing microglia in the midbrain of *rraga* mutants at 3 dpf, but not in earlier stages (Fig. 1D). To investigate how the reduction of macrophage numbers in the brain occurs in *rraga* mutants, we performed timelapse imaging starting at 60 hpf and ending at ∼78 hpf. Analysis of the timelapse movies showed that fewer macrophages migrate into the brain in *rraga* mutants (Fig. 1E-H, Movies S1 and S2). Furthermore, timelapse imaging provided no evidence for other possibilities, such as reduced proliferation or increased apoptosis in *rraga* mutants (Fig. 1E-H, Movies S1 and S2) after embryonic macrophages enter the brain. These experiments demonstrate that microglial progenitors in *rraga* mutants fail to migrate into the brain between 2 and 3 dpf.

### Peripheral macrophages are disrupted in *rraga* mutants

Previous studies have not examined the extent to which macrophages outside the brain are affected in *rraga* mutants. We used the *Tg(mpeg:GFP)* transgene to visualize macrophages at 2, 2.5, and 3 dpf. In lateral views of the head, the numbers of macrophages were similar between *rraga* mutants and their wildtype siblings at 2 dpf (Fig. 1I, 1K, fig. S1E), and *rraga* mutants showed a slight, but insignificant, reduction of macrophages in this region at 2.5 dpf (Fig. 1K, fig. S1A, S1E). There was, however, a striking and significant reduction of macrophage numbers in the head in *rraga* mutants by 3 dpf (Fig. 1I’, 1K, fig. S1E), with very few macrophages remaining in the mutants. Although macrophage numbers were reduced in *rraga* mutants beginning at 2.5 dpf in the trunk and tail regions (Fig. 1J-J’, 1L, fig. S1A and S1E), many macrophages persisted in outside the head in *rraga* mutants at all stages we examined up to 8 dpf (Fig. 1J-J’, 1L, fig. S1B-D). Next, to determine if the reduced macrophage numbers in *rraga* mutants results from reduced proliferation of these cells, we used in vivo EdU labeling by pulsing the animals with EdU at 2.5 dpf and chasing up to 3 dpf. We observed an insignificant decrease in the number of proliferating macrophages in *rraga* mutants (Fig. 1M). However, we cannot rule out subtle effects in proliferation, apoptosis, or specification that could affect peripheral macrophage cell numbers. An interesting feature of macrophages and microglia is that the morphology of these cells is frequently used as a readout of their functional state; highly ramified microglia are typically thought to be “resting” or “surveilling”, whereas rounded microglia are considered to be activated in response to infection or injury, or are dysfunctional in some way (*45*). Starting at 3 dpf, macrophages in the tail region of *rraga* mutants displayed an abnormal, rounded morphology (Fig. 1N), similar to microglia in these mutants (*41*). Together, these observations show that the number, migration, and morphology of macrophages are disrupted in *rraga* mutants and that these phenotypes manifest between 2 and 3 dpf.

### Macrophages in *rraga* mutants show a significant upregulation of lysosomal pathways and downregulation of immune and migration-related genes

To further investigate the molecular mechanisms underlying the defects in the macrophage lineage in *rraga* mutants, we performed RNA-Sequencing on macrophages and microglia isolated from these mutants. We crossed *rraga* heterozygotes carrying the *mpeg:GFP* transgene, scored the mutant and wildtype animals using confocal microscopy, dissociated the larvae, sorted GFP^+^ cells at 6 dpf, extracted RNA, and performed RNA-Sequencing (Fig. 2A). Gene Ontology term enrichment analysis of the genes upregulated in macrophages from *rraga* mutants revealed a strong enrichment of the terms Vacuole, Lysosome, Lytic Vacuole, and Endosome (Fig. 2B, 2C, fig. S2A, Table S2). To validate these transcriptomic observations, we generated genetic tools and used chemical probes to assess the endolysosomes in macrophages and microglia in vivo. LAMP1 and LAMP2 are estimated to contribute 50% of all proteins in lysosomal membrane (*46*), and these genes were significantly upregulated in our dataset (Fig. 2B). We generated zebrafish LAMP1-mCherry and LAMP2-mCherry constructs and used the *mpeg* promoter to express these transgenes in the macrophage lineage (Fig. 2D). In parallel, we also generated mCherry-Rab5 or mCherry-Rab7 constructs, driven under the *mpeg* promoter, to label early and late endosomes respectively (*47*). Consistent with our RNA-Seq data, we found a significant expansion of LAMP1-mCherry, LAMP2-mCherry, mCherry-Rab5, and mCherry-Rab7 punctae in the microglia of *rraga* mutants relative to their wildtype siblings (Fig. 2D, 2E).

**FIG. 2:**
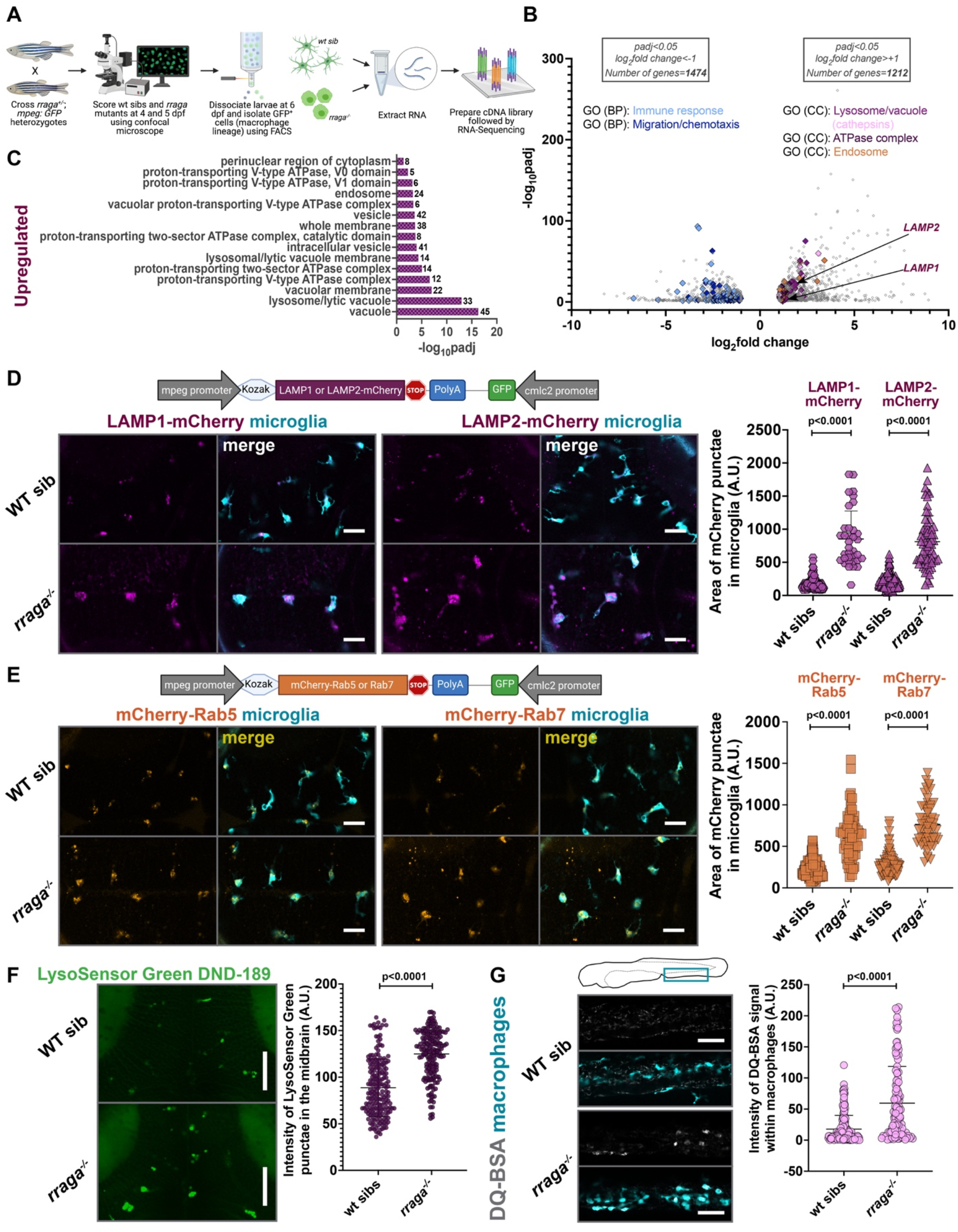
Lysosomal genes are significantly upregulated and endolysosomes are expanded in macrophages isolated from *rraga* mutants. (A) Experimental schematic for RNA-Sequencing. (B) Volcano plot showing significantly differentially upregulated and downregulated genes in the macrophages of *rraga* mutants relative to wildtype siblings. (C) Gene Ontology (Cellular Component) enrichment analysis showing an upregulation of lysosome-associated terms in macrophages from *rraga* mutants. (D) Imaging and quantitation of LAMP1 or LAMP2-mCherry transgene expression in *rraga* mutants and wildtype siblings at 4 dpf. Scale bars, 20 µm. (E) Imaging and quantitation of mCherry-Rab5 or Rab7 transgene expression in *rraga* mutants and wildtype siblings at 4 dpf. Scale bars, 20 µm. (F) LysoSensor Green labeling and quantitation of LysoSensor Green intensity at 4 dpf. Scale bars, 50 µm. (D-F) graphs show mean with standard deviation; significance was determined using unpaired t-test. (G) DQ-BSA labeling at 4 dpf and quantitation of DQ-BSA intensity. Scale bars, 50 µm. Graph shows mean with standard deviation; significance was determined using Mann-Whitney U test.

Our analysis also revealed a significant expansion of proton-transporting vATPase complex genes (Fig. 2B, 2C, fig. S2A), which are central players in intracellular acidification and organelle pH control (*48*). To test intracellular acidification of microglia, we treated *rraga* mutant larvae with the dye LysoSensor Green DND-189, which becomes more fluorescent in acidic compartments of the cell (*49*). As suggested by our RNA-Seq data, we found a significant increase of the LysoSensor Green intensity in apparent microglia in the midbrain of *rraga* mutants relative to their wildtype siblings (Fig. 2F). Finally, several cathepsin genes, which encode lysosomal proteases activated by acidic pH, were upregulated in *rraga* mutants (*50*) (Fig. 2B, Table S2). To assay this apparent increase in proteolytic activity in the macrophages of *rraga* mutants, we used DQ-BSA (*51*) and Magic Red (MR) Cathepsin (*52*), dyes that are non-fluorescent in neutral pH but fluoresce brightly when proteases cleave to release the quenching residues in these dyes. Once again, in accord with our RNA-Seq data, we found fluorescence of both DQ-BSA and MR-Cathepsin increased in macrophages of *rraga* mutants (Fig. 2G, fig. S2B). Our results demonstrate that endolysosomal gene expression is expanded and acid-activated protease activity is increased in macrophages of *rraga* mutants in vivo.

The gene enrichment terms most significantly downregulated in macrophages from *rraga* mutants were immune response and migration/taxis, followed by lipid metabolism and other metabolic pathways (Fig. 2B, 3A, 3E, fig. S2C, S2D, Table S3). To examine possible abnormalities in macrophage clearance of microbial debris, we challenged the tail macrophages of *rraga* larvae with *E. coli* particles labeled with Texas Red. There was a dramatic reduction in the uptake of these *E. coli* particles by *rraga* mutants relative to their wildtype siblings (Fig. 3B, fig. S2E). In contrast, macrophages in *rraga* mutants can sense and ingest weakly immunogenic molecules such as Dextran Texas Red and Ovalbumin 555 (Fig. 3C, 3D), suggesting that RagA is differentially required for engulfment or processing of different substrates. The downregulation of genes corresponding to migration and taxis in macrophages from *rraga* mutants (Fig. 3E) corroborates our earlier observation that early macrophages do not migrate into the brain of *rraga* mutants to become microglia (Fig. 1E-H). To further assess macrophage migration in *rraga* mutants, we used timelapse imaging to track macrophage movement in response to injury of the tail fin (Fig. 3F). Significantly fewer macrophages in *rraga* mutants moved to the wound site at 8 hours post injury (Fig. 3F). When combined with our RNA-Seq analysis, the abnormalities in macrophage migration during brain colonization (Fig. 1E-H) and wound response (Fig. 3F) indicate that cells of the macrophage lineage in *rraga* mutants are unable to sense or respond appropriately to developmental signals (*35, 53*) or injury cues. Together, our RNA-Seq data illuminate heretofore unappreciated functions of RagA in the macrophage lineage, particularly the contribution of RagA in activation of innate immune response and cell migration.

**FIG. 3:**
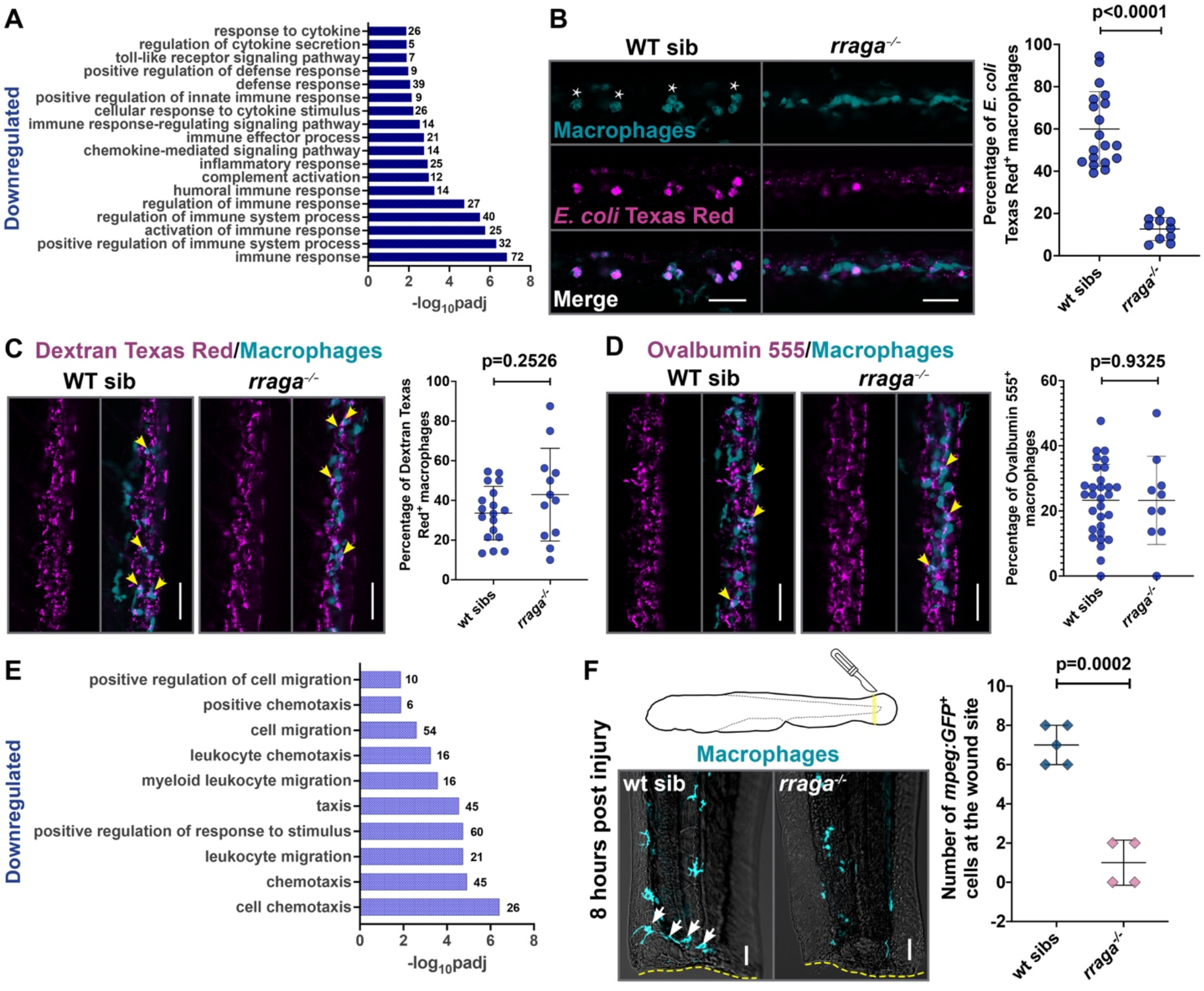
Macrophages show defective microbial clearance and migration in *rraga* mutants. (A) Bar graph showing GO terms (Biological Process) related to immune system functions that are significantly downregulated in macrophages from *rraga* mutants. Injection of (B) *E. coli* Texas Red, (C) Dextran Texas Red, (D) Ovalbumin 555 to challenge the macrophages in *rraga* mutants and siblings at 4 dpf and corresponding quantitation. Arrows in (C) and (D) denote colocalization of debris with macrophages. Note that wildtype macrophages responding to *E. coli* become activated and display amoeboid morphology (marked with asterisks). Scale bars, 50 µm. All graphs show mean with standard deviation; significance was determined using Mann-Whitney U test. (E) GO terms related to migration/taxis that are significantly downregulated in macrophages from *rraga* mutants. (F) Macrophage response to tail injury at 4 dpf. Arrows in the wildtype image show macrophages at the wound site (yellow dotted line). Scale bars, 20 µm. Graph shows mean with standard deviation; significance was determined using unpaired t-test.

### Simultaneous loss of lysosomal transcription factors *tfeb, tfe3a*, and *tfe3b* rescues microglia numbers in *rraga* mutants

The transcription factor EB (Tfeb) activates most lysosomal functions including lysosomal biogenesis, autophagy, and exocytosis (*15-17*). Several lines of evidence indicate that RagA represses Tfeb and other related transcription factors (*14, 18*), although the extent to which the RagA-Tfeb lysosomal pathway functions in the macrophage lineage in vivo is unclear. To address whether RagA represses Tfeb in microglia, we crossed *rraga*^*+/-*^; *tfeb*^*+/-*^ double heterozygotes (*54*), and labeled microglia using neutral red, a dye that preferentially accumulates in microglia (*44*). Imaging and quantification of microglia in the dorsal midbrain showed that *rraga* mutants have significantly fewer neutral red-labelled microglia, as shown previously (*41*) (Fig. 4A). Loss of *tfeb* in *rraga* mutants partially restored microglia numbers (Fig. 4A, B). These data provided evidence that hyperactivated Tfeb contributes to, but does not solely cause, microglial defects in *rraga* mutants.

**FIG. 4:**
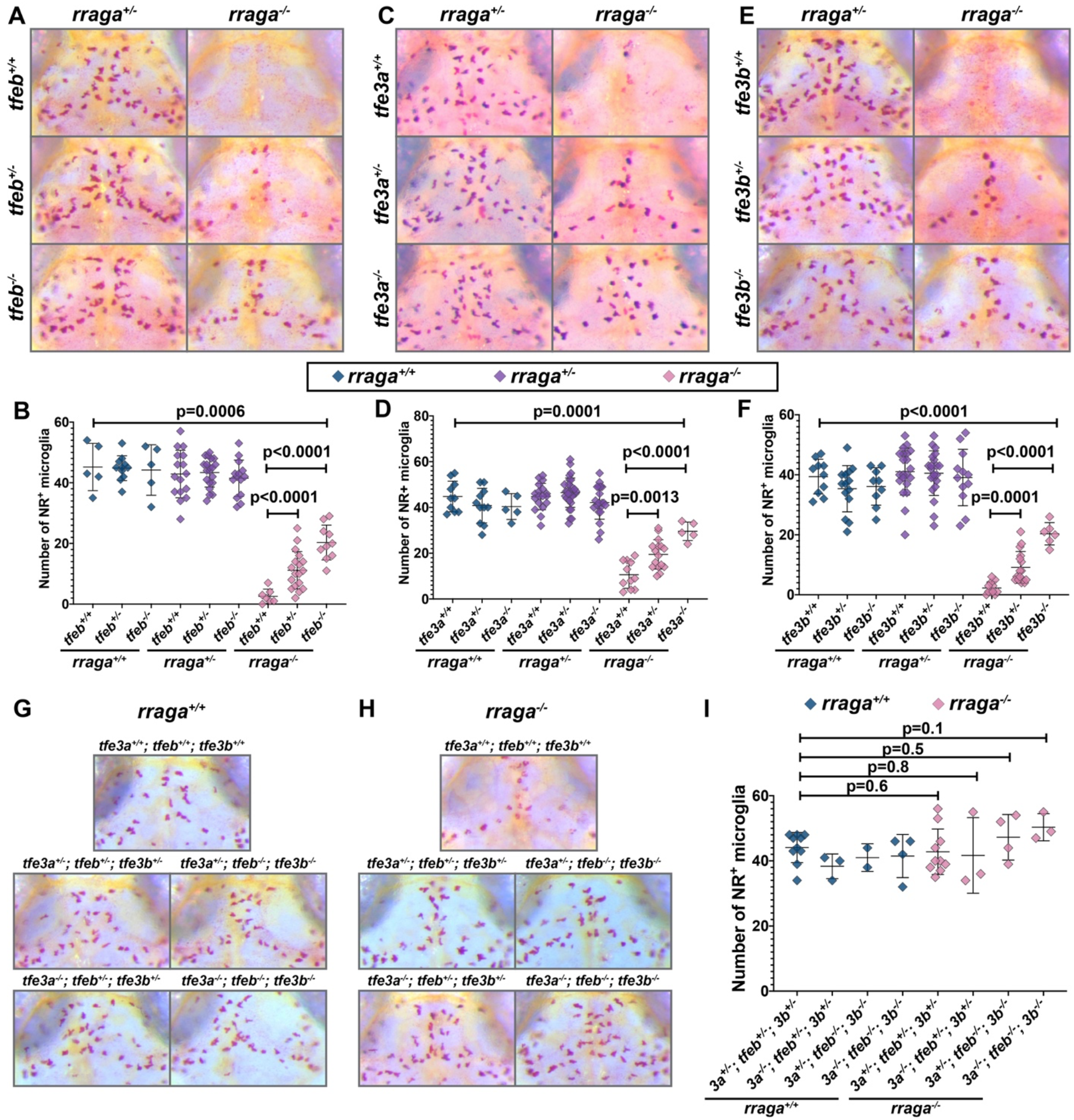
Microglia numbers are restored in *rraga* mutants with simultaneous loss of *tfeb, tfe3a*, or *tfe3b*. Images show dorsal views of the midbrain (anterior on top) of neutral red (NR) stained larvae of indicated genotypes at 5 dpf and graphs show corresponding quantification. (A, B) progeny from *rraga*^*+/-*^; *tfeb*^*+/-*^ intercross; (C, D) progeny from *rraga*^*+/-*^; *tfe3a*^*+/-*^ intercross; (E, F) progeny from *rraga*^*+/-*^; *tfe3b*^*+/-*^ intercross; and (G, H, I) progeny *rraga*^*+/-*^; *tfeb*^*+/-*^; *tfe3a*^*+/-*^; *tfe3b*^*+/-*^ quadruple heterozygous intercross. Images of a *rraga* mutant and *tfeb* and *tfe3* genotypes that result in complete rescue of NR^+^ microglia in *rraga* mutants are shown in (H). In (G), the same genotypes, but in a background wildtype for *rraga*, are included for comparison. All graphs show mean with standard deviation; significance was determined using unpaired t-test.

One hypothesis for the incomplete rescue we observed in *rraga; tfeb* double mutants is that RagA might also repress other members of the Microphthalmia-associated transcription factor (MiTF) family, which consists of Mitf, Tfeb, Tfe3, and Tfec (*14*). Notably, Tfeb and Tfe3 have been shown to act cooperatively to regulate cytokine production in the RAW 264.7 macrophage cell line (*55*), although the extent of functional overlap between these transcription factors in macrophages in vivo remains uncharacterized. To determine if Tfe3 functions in macrophages and microglia, we generated mutants for the two zebrafish paralogs of *tfe3, tfe3a* and *tfe3b* (fig. S3 and Table S1) (*56*). Neutral red assay revealed that mutations in *tfe3a* (Fig. 4C, 4D) or *tfe3b* (Fig. 4E, 4F) can partially rescue microglia numbers in *rraga* mutants, much as observed in *rraga; tfeb* double mutants (Fig. 4A, 4B). Importantly, the loss of even a single copy of *tfeb, tfe3a*, or *tfe3b* can partially rescue microglia in *rraga* mutants (*rraga*^*+/-*^; *tfeb*^*+/-*^ in Fig. 4B; *rraga*^*+/-*^; *tfe3a*^*+/-*^ in Fig. 4D; *rraga*^*+/-*^; *tfe3b*^*+/-*^ in Fig. 4F), indicating that microglia are highly sensitive to increased levels of each of these transcription factors.

To further explore the dosage-dependent rescue of Tfeb and Tfe3, as well as to better understand the extent of functional overlap between Tfeb, Tfe3a, and Tfe3b, we generated *rraga*^*+/-*^; *tfeb*^*+/-*^; *tfe3a*^*+/-*^; *tfe3b*^*+/-*^ quadruple heterozygous animals. Neutral red assay and quantification (Fig. 4G-I) revealed that combined loss of a single copy of *tfeb, tfe3a*, and *tfe3b* was sufficient to completely restore microglia numbers in *rraga* mutants (Fig. 4H). We also observed a full rescue of microglia in *rraga* mutants when 4, 5, or 6 copies of *tfeb, tfe3a*, and *tfe3b* were mutated (Fig. 4H, 4I). We did not observe any obvious defects in microglia in *tfeb, tfe3a*, or *tfe3b* mutants alone (Fig. 4A-F), or in combination, including in *tfeb; tfe3a; tfe3b* triple mutants (Fig. 4G-I). Collectively, these observations indicate that RagA represses Tfeb, Tfe3a, and Tfe3b in microglia, that hyperactivity of these transcription factors causes the microglia defects in *rraga* mutants, and that *tfeb* and its homologs are not required for normal microglial development.

### *folliculin* and *rraga* mutants have similar phenotypes

Several studies link the protein Folliculin (Flcn) to the repression of Tfe3 (*57-59*), and in some cases, Tfeb (*60*); however, the mechanism through which Flcn regulates the activation of Tfe3 (or Tfeb) is not fully understood. To assess the functional interaction between Flcn and Tfeb/Tfe3, we generated *flcn* mutants using CRISPR-Cas9 (fig. S4A). We found that mutations in *flcn* or *rraga* caused a strikingly similar phenotype in macrophages and microglia. *flcn* mutants have significantly fewer neutral red-labeled microglia in their dorsal midbrain (Fig. 5A, fig. S4B), as in the case of *rraga* mutants (Fig. 4) (*41*). Furthermore, this reduction in microglia numbers in *flcn* mutants was rescued by simultaneous loss of *tfeb* and *tfe3b* (Fig. 5A) and partially rescued in *flcn; tfe3a* double mutants (fig. S4B). Once again, as in the case of *rraga* mutants (Fig. 4), we observed a dosage-dependent effect, and *flcn* mutants that are heterozygous for the *tfeb; tfe3b* double mutant chromosome or *tfe3a* display a partial, intermediate level of rescue of microglia number in the neutral red assay (Fig. 5A, fig. S4B).

**FIG. 5:**
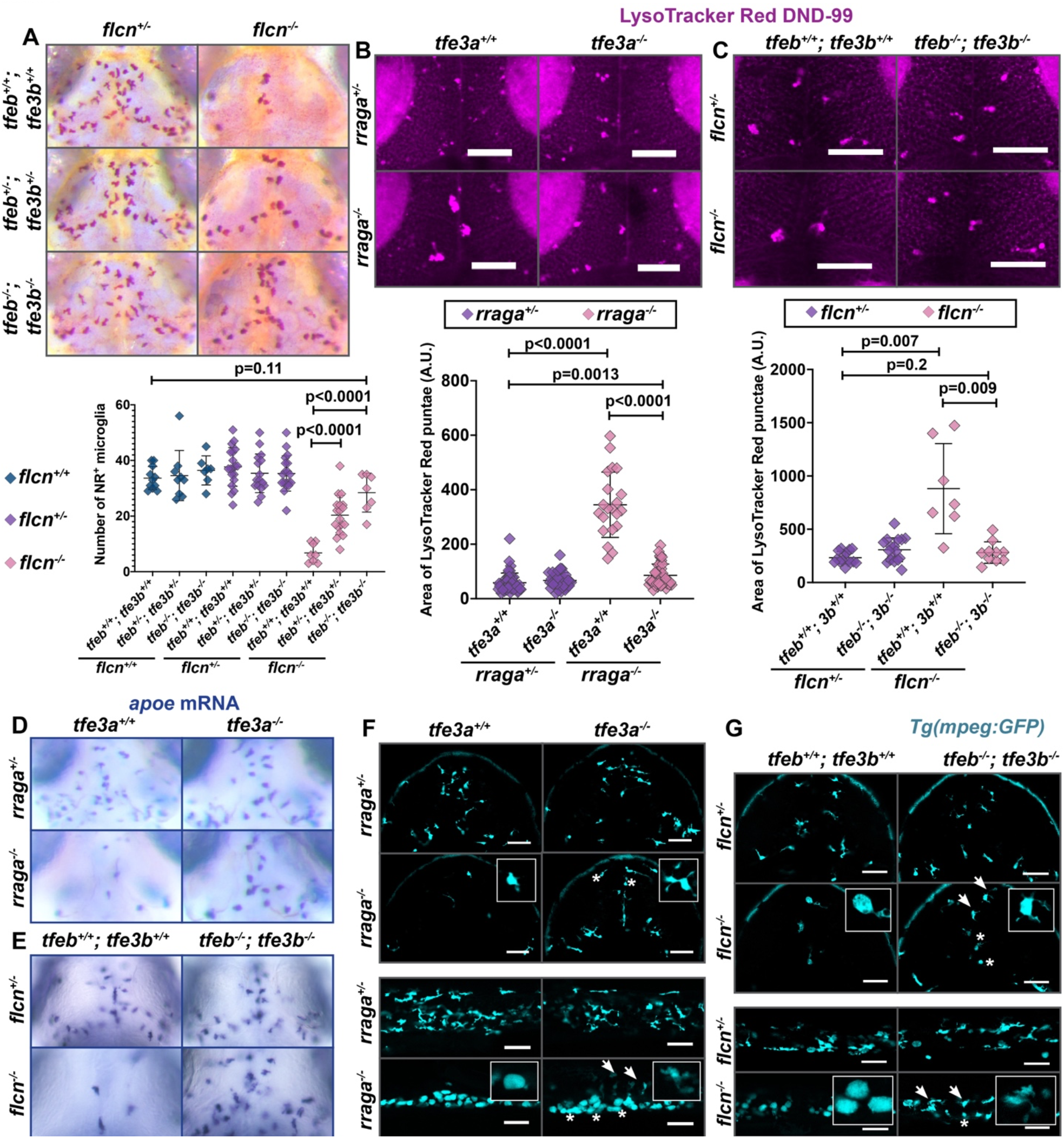
Simultaneous mutation in *tfeb, tfe3a*, or *tfe3b* rescues *flcn* and *rraga* mutant phenotypes. (A) NR assay and quantification of microglia in larvae obtained from *flcn*^*+/-*^; *tfeb*^*+/-*^; *tfe3b*^*+/-*^ intercross. (B, C) LysoTracker Red assay and quantification of the area of LysoTracker Red punctae in microglia of larvae from (B) *rraga*^*+/-*^; *tfe3a*^*+/-*^ intercross and (C) *flcn*^*+/-*^; *tfeb*^*+/-*^; *tfe3b*^*+/-*^ intercross. Images show LysoTracker Red signal in dorsal view of the midbrain. (D, E) *apoe* mRNA expression at 4 dpf and rescue of microglia in the progeny of (D) *rraga*^*+/-*^; *tfe3a*^*+/-*^ intercross, and (e) *flcn*^*+/-*^; *tfeb*^*+/-*^; *tfe3b*^*+/-*^ intercross. (F, G) *mpeg:GFP* expression to visualize microglia and macrophages in (F) *rraga; tfe3a* double mutants and (G) *flcn; tfeb; tfe3b* triple mutants. Arrows denote microglia and macrophages in which ramified morphology has been restored, asterisks denote cells with amoeboid morphology, and insets show magnified views of cell morphology. Scale bars, 50 µm. Graphs show mean with standard deviation; significance was determined using unpaired t-test.

We also used the dye LysoTracker Red (*36*) to examine the morphology of acidic compartments within microglia in *rraga* or *flcn* mutants. As shown previously (*41*), *rraga* mutants have a significant expansion of LysoTracker Red punctae in microglia (Fig. 5B); we found that this expansion is partially rescued in *rraga; tfe3a* double mutants (Fig. 5B). Remarkably similar to *rraga* mutants, we also found that *flcn* mutants had enlarged LysoTracker Red punctae in their microglia and that this expansion is fully rescued in *flcn; tfeb; tfe3b* mutants (Fig. 5C). *rraga* mutants have very few *apoe-*expressing microglia in their dorsal midbrain (*41*) (Fig. 5D), and we found that this defect is partially rescued in *rraga; tfe3a* double mutants (Fig. 5D). Similarly, there was a strong reduction in *apoe*-expressing microglia in *flcn* mutants and this defect was rescued in *flcn; tfeb; tfe3b* mutants (Fig. 5E). Finally, we examined these genotypes using the *Tg(mpeg:GFP)* line to assess the morphology of microglia and macrophages. Both *rraga* and *flcn* mutants displayed abnormal, rounded morphology of microglia and macrophages (Fig. 5F, 5G), a defect that was at least partially restored in *rraga; tfe3a* mutants (Fig. 5F) and *flcn; tfeb; tfe3b* mutants (Fig. 5G). In sum, our experiments show that the key lysosomal proteins Flcn and RagA have very similar functions in microglia and macrophages in vivo and are both essential to repress Tfeb and Tfe3. Moreover, all the macrophage number and morphology defects observed in *rraga* or *flcn* mutants can be rescued to varying degrees by simultaneous loss of *tfeb*, or *tfe3a*, or *tfe3b*.

### Overexpression of *tfe3b* in macrophages recapitulates *rraga* mutant phenotypes in microglia and macrophages

We reasoned that if Tfeb and Tfe3 are the primary downstream targets repressed by RagA and Flcn, then overexpression of Tfeb or Tfe3 in macrophages should recapitulate the *rraga* or *flcn* mutant phenotypes. We constructed transgenic fish that overexpressed Tfe3b in cells of the macrophage lineage, *Tg(mpeg:tfe3b; cmlc2:mCherry)*, and found that indeed these animals had the defects characteristic of *rraga* or *flcn* mutants (Fig. 6A-E). First, *Tg(mpeg:tfe3b)* animals have significantly fewer neutral red-labeled microglia than *Tg(mpeg:GFP)* animals (Fig. 6A), similar to mutants for *rraga* or *flcn* (Fig. 4, 5A, fig. S4B) (*41*). Second, animals that overexpress *tfe3b* in macrophages have significantly enlarged LysoTracker Red punctae in microglia (Fig. 6B), as observed in *rraga* or *flcn* mutants (Fig. 5B, 5C). Third, *Tg(mpeg:tfe3b)* animals had reduced *apoe* labeling in the dorsal midbrain (Fig. 6C), once again similar to both *rraga* and *flcn* mutants (Fig. 5D, 5E) (*41*), and in contrast to *Tg(mpeg:GFP)* controls (Fig. 6C). Finally, by crossing *Tg(mpeg:tfe3b)* animals to the *Tg(mpeg:GFP)* fish, we found that animals overexpressing *tfe3b* in the macrophage lineage had fewer microglia and macrophages, and that the remaining cells exhibited abnormal, rounded morphology (Fig. 6D, 6E), similar to *rraga* (Fig. 1N, 5F) (*41*) or *flcn* (Fig. 5G) mutants. These experiments demonstrate that the overexpression of *tfe3b* is sufficient to recapitulate the *rraga* mutant phenotype in microglia and macrophages (Fig. 6A-E). Collectively, our data indicate that the primary essential function of RagA and Flcn in macrophages is to repress Tfeb and Tfe3.

**FIG. 6:**
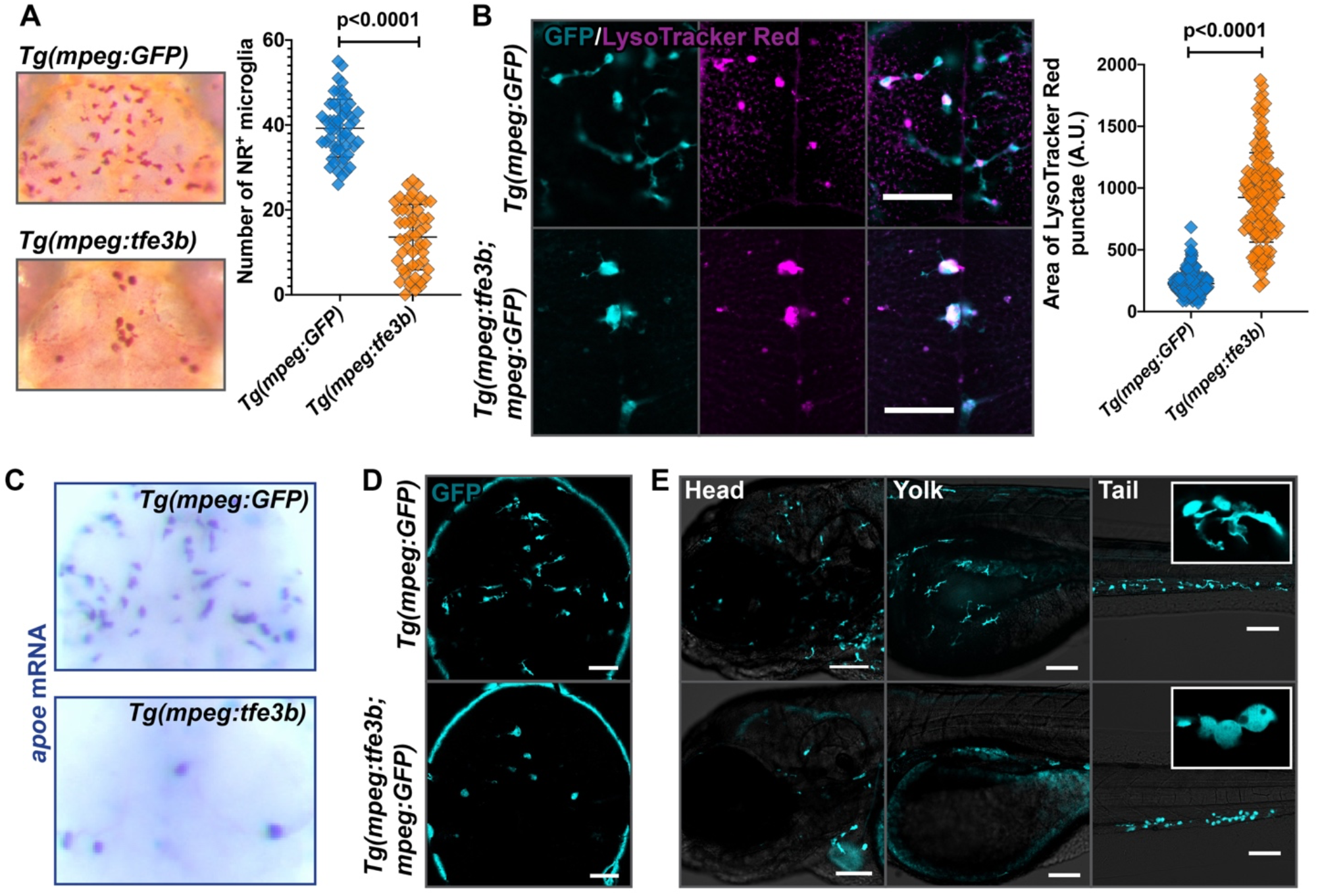
Overexpression of *tfe3b* in the macrophage lineage disrupts microglia number and morphology as in *rraga* mutants. Comparison of macrophages and microglia between animals overexpressing Tfe3b in the macrophage lineage, *Tg(mpeg:tfe3b)*, and controls, *Tg(mpeg:GFP)*. (A) Neutral red assay and quantification, (B) LysoTracker Red assay and quantification, (C) *apoe* in situ hybridization, and (D, E) live imaging with the *mpeg:GFP* transgene in (D) the brain and (E) head, yolk, and tail regions. Insets show magnified views of cell morphology. Scale bars, 50 µm. All graphs show mean with standard deviation; significance was determined using unpaired t-test.

### Tfeb and Tfe3 activate lysosomal pathways under conditions of cellular stress but are dispensable for basal expression of these genes

To test the requirement of Tfeb or Tfe3 for the specification, maintenance, or function of microglia and macrophages, we generated *tfeb; tfe3a; tfe3b* triple mutants. Corroborating our earlier analysis, microglia and macrophages in the triple mutants are indistinguishable from their wildtype counterparts in both number and morphology (Fig. 4, 7A, fig. S5A). To examine whether macrophages in the triple mutants respond differently from their wildtype siblings in response to environmental challenges, we injured the tail fins of these animals and found that macrophages in triple mutants respond to tail injury similarly to their siblings (Fig. 7B). Additionally, we injected Zymosan A into the circulation of these larvae and found once again that activation of macrophages in triple mutants is comparable to that of their wildtype siblings (amoeboid morphology in Fig. 7C). These observations indicate that Tfeb, Tfe3a, and Tfe3b are not necessary for the survival of microglia and macrophages or for their response to injury or inflammatory stimulus.

**FIG. 7:**
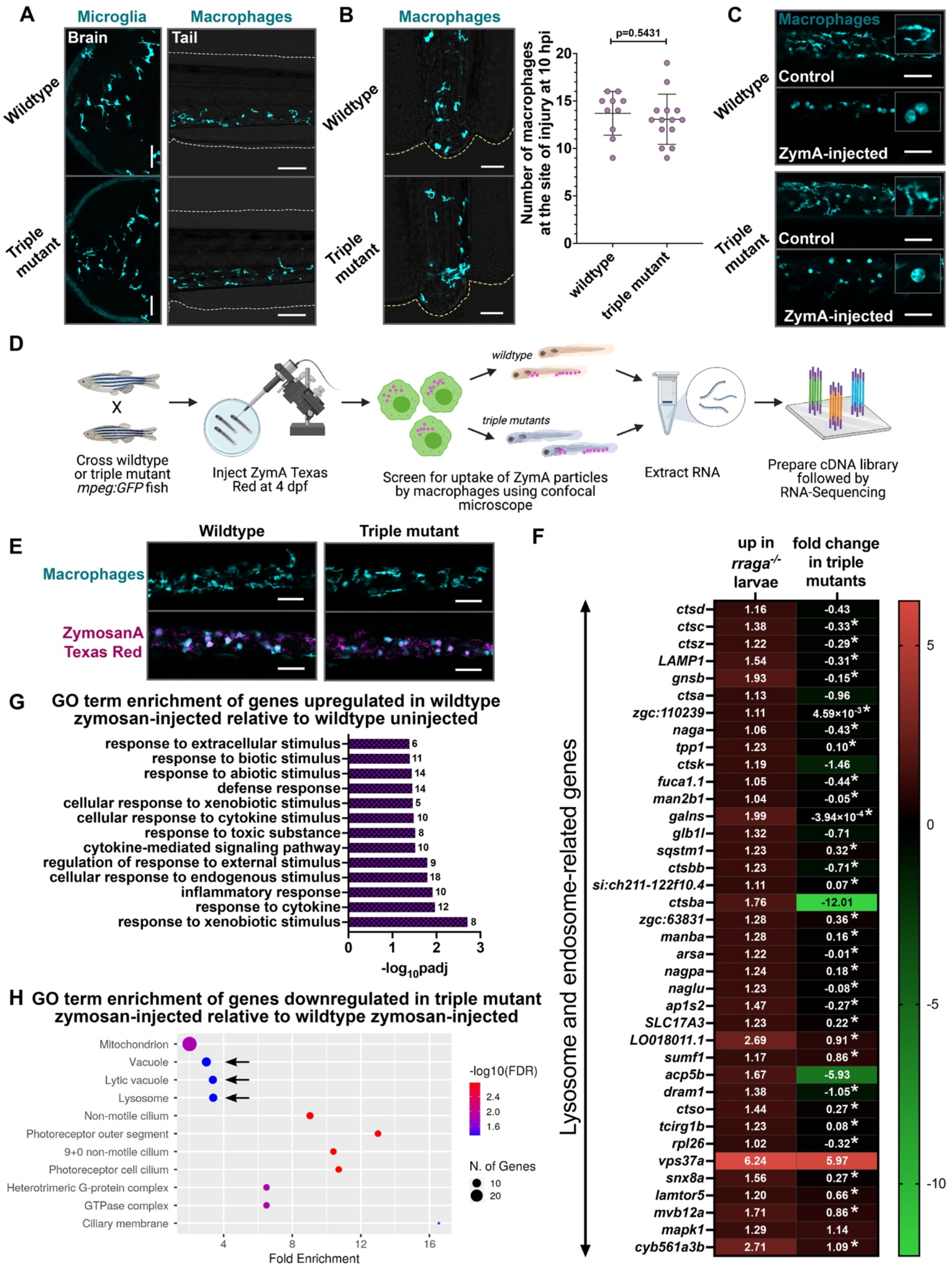
Tfeb and Tfe3 activate lysosomal pathways only under conditions of stress. (A) Microglia and macrophages visualized using the *mpeg:GFP* transgene in wildtype animals and *tfeb; tfe3a; tfe3b* triple mutants at 4 dpf. Macrophage response to (B) tail injury and (C) systemic Zymosan A injection in wildtype and triple mutant animals at 4 dpf. Insets in (C) show magnified views of macrophage morphology. Graph shows mean with standard deviation; significance was determined using unpaired t-test. (D) Experimental schematic for RNA-Sequencing. (E) Microscopy-based validation of Zymosan A Texas Red uptake by the trunk and tail macrophages in triple mutants and wildtype animals at 4 dpf. (F) Heat map depicting all the lysosomal genes significantly upregulated (log_2_fold>+1, padj<0.05) in *rraga* mutant whole larvae and the corresponding fold change of the gene in triple mutant larval RNA preparations. The log_2_fold change is shown in each cell; fold change values with asterisks are not significant (padj>0.05). (G) GO term-enrichment analysis of genes differentially upregulated in wildtype Zymosan A-injected animals relative to uninjected wildtype controls. (H) GO term-enrichment of genes significantly downregulated in Zymosan A-injected triple mutants.

Activation of Tfeb or Tfe3 upregulates lysosomal genes responsible for lysosomal biogenesis, autophagy, and exocytosis in cultured cells (*14*). Despite the sufficiency of Tfeb and Tfe3 to activate lysosomal pathways, the extent to which these transcription factors are *necessary* for the expression of lysosomal genes is unknown. Furthermore, there is evidence that Tfeb and Tfe3 mediate cellular stress response (*61, 62*), but the extent to which they drive the macrophage response to various environmental challenges remains unclear. In light of our surprising observation that the *tfeb; tfe3a; tfe3b* triple mutants are viable, with normal appearing microglia and macrophages, we decided to perform RNA-Sequencing on whole larvae to elucidate the functions of Tfeb and Tfe3. Briefly, we crossed wildtype or triple mutant fish carrying the *mpeg:GFP* transgene and injected Zymosan A Texas Red into the yolk of some of these larvae at 4 dpf (Fig. 7D). We screened for the uptake of ZymA particles by macrophages using confocal microscopy (Fig. 7D, 7E), pooled ZymA-injected and uninjected animals of wildtype and triple mutant genotypes separately, extracted RNA, prepared cDNA libraries, and performed RNA-Sequencing (Fig. 7D).

Comparison of the *tfeb; tfe3b; tfe3a* triple mutants to wildtype animals yielded 366 upregulated (log_2_fold>+1; padj<0.05) and 511 downregulated genes (log_2_fold<-1; padj<0.05), but there was no GO term enrichment in genes differentially up or downregulated in triple mutants (Table S4). We also performed whole larvae RNASeq on *rraga* mutants and their wildtype siblings to determine the extent to which lysosomal genes are enriched in whole larvae preparations of *rraga* mutants as well as to compare the differential lysosomal gene expression between *rraga* and *tfeb; tfe3a; tfe3b* mutants. As expected, *rraga* mutants show a significant upregulation of lysosomal genes; however, expression of most of these genes did not change significantly in the triple mutants, with some exceptions (Fig. 7F). These results revealed that Tfeb and Tfe3 are not necessary for expression of lysosomal pathways in whole animals under normal conditions. Comparison of genes differentially upregulated in the ZymA-injected versus uninjected wildtype animals revealed expected enrichment of GO categories corresponding to defense response, response to various kinds of stimuli, as well as an inflammatory signature (Fig. 7G). Notably, GO categories Vacuole, Lytic Vacuole, and Lysosome were significantly downregulated in ZymA-injected triple mutants versus ZymA-injected wildtype animals (Fig. 7H). These terms are not significantly enriched in the corresponding dataset of genes downregulated in uninjected triple mutants versus uninjected wildtype animals. In another enrichment approach, KEGG pathway analysis revealed enrichment of the category Lysosome in the genes downregulated in ZymA-injected triple mutants, while this category is not significantly changed in the uninjected triple mutants (fig. S5B, S5C). Finally, Ingenuity Pathway Analysis yielded phagosome maturation and CLEAR signaling pathway (lysosomal genes activated by Tfeb and Tfe3 (*15*)) as top canonical pathways when we analyzed the genes downregulated in ZymA-injected triple mutants (fig. S5D). Together, our results indicate that Tfeb and Tfe3 are dispensable for basal levels of lysosomal gene expression under homeostatic conditions in vivo, but these transcription factors are required for activation of lysosomal pathways in response to cellular stress.

## Discussion

Our experiments define an antagonistic regulatory circuit – involving RagA and Flcn on one side, and Tfeb, Tfe3a, and Tfeb on the other – that is essential for the development and function of macrophages in vivo. Using transcriptomic analyses, we demonstrate that macrophages from *rraga* mutants display a strong upregulation of known lysosomal target genes of Tfeb and Tfe3; correspondingly, we observe a significant expansion of the endolysosomal compartment in these mutants in vivo. Simultaneously, we uncover a previously unappreciated role for RagA in the regulation of genes involved in migration and innate immunity, and our cellular studies reveal disrupted migration of macrophages and abnormal processing of microbial substrates by these cells in *rraga* mutants. Our genetic studies show that loss of Tfeb and Tfe3 can rescue microglia in *rraga* and *flcn* mutants, and macrophages overexpressing Tfe3b phenotypically resemble those in *rraga* mutants. Thus, our experiments demonstrate that the central function of RagA and Flcn in macrophages is to repress Tfeb and Tfe3.

Notably, our experiments reveal that cells of the macrophage lineage are acutely sensitive to the levels of Tfeb, Tfe3, and Tfe3b, because loss of even a single copy of one of these three genes is sufficient to partially restore the microglia numbers in *rraga* or *flcn* mutants. Overexpression of Tfe3b alone is sufficient to phenocopy the microglia number and morphology defects seen in *rraga* and *flcn* mutants, indicating that Tfeb, Tfe3a, and Tfe3b are at least partly functionally redundant in the macrophage lineage. Interesting future directions will be to investigate the extent to which the targets of the MiTF family of transcription factors overlap in different cell types, and to uncover the mechanisms by which cells sense the activity levels of these transcription factors.

Microglia and macrophages depend on lysosomal pathways to execute their phagocytic functions; it is therefore seemingly paradoxical that microglia and macrophages appear normal in *tfeb; tfe3a; tfe3b* triple mutants. Indeed, *tfeb; tfe3a; tfe3b* triple mutant animals are viable and fertile as adults, although they seem to grow at a slower rate than their wildtype siblings. Our whole larvae transcriptomic studies revealed that Tfeb, Tfe3a, and Tfe3b are not required for basal expression of lysosomal and autophagy genes, indicating that other pathways must control lysosome biogenesis and activity during normal development. We show that Tfeb and Tfe3 are required to activate lysosomal pathways in whole larvae under conditions of stress, providing compelling in vivo evidence for the role of these transcription factors as mediators of the cellular stress response (*61-63*). A more detailed analysis of Tfeb and Tfe3 targets under different conditions of stress, such as apoptotic debris, misfolded proteins, or microbial infection, may reveal the entire range of lysosomal and other pathways activated by these transcription factors in response to specific environmental challenges. Future studies will also address how different cell types employ RagA, Flcn, Tfeb, and Tfeb3 to detect and respond to diverse range of stressors.

Finally, attempts to enhance Tfeb-mediated lysosomal pathways in animal models of neurodegenerative diseases (*19, 20, 22, 23, 64*), particularly Alzheimer’s disease (AD) (*27-29*), have led to beneficial effects, but the identity of cells that must overexpress Tfeb in vivo to generate these favorable cognitive outcomes remains unclear. Since prolonged activation of inflammatory pathways in AD results in neurotoxicity and indiscriminate loss of viable cells leading to cognitive defects (*65-69*), our observation that hyperactivation of Tfeb and Tfe3 may downregulate immune pathways in the macrophage lineage may offer a key insight into the beneficial outcomes observed in animal models of AD. Moreover, while we find that repression of Tfeb and Tfe3 is necessary for the development and migration of embryonic macrophages, studying the role of these transcription factors in the maintenance and function of adult microglia presents an exciting future direction (*38, 70, 71*). Future studies aiming to understand the regulation and functions of Tfeb and Tfe3 in microglia during development, disease, and aging will not only broaden our understanding of lysosomal pathways in microglia, but will also inform therapeutic strategies aimed at modulating Tfeb activity in neurodegeneration and other diseases.

## Materials and Methods

### Zebrafish lines

Embryos from wildtype (TL or AB) strains and *Tg(mpeg:GFP)* transgenic line (*42*) were raised at 28.5 ºC in embryo water with methylene blue. For all imaging experiments, embryos were treated with 0.003% 1-phenyl-2-thiourea (PTU) to inhibit pigmentation between 10 and 24 hours post fertilization, and anesthetization was done using .016% MS-222 (Tricaine) prior to experimental procedures. For neutral red assay, methylene blue was excluded from embryo water. All animal protocols have been approved by the Stanford Institutional Animal Care and Use Committee.

### Timecourse and timelapse experiments

Embryos for all timecourse and timelapse experiments were staged between 0 and 12 hpf and typically dechorionated between 30-36 hpf. For the 24-26 hpf timepoint, embryos were dechorionated immediately prior to mounting. Genotyping was typically performed after image acquisition using Zeiss LSM confocal microscope. For all timecourse experiments, cells were first counted, and a single, high-resolution, representative plane was subsequently imaged using the 20X objective.

For the timelapse experiments starting at 60 hpf, a region of interest in the brain containing the midbrain, forebrain and a portion of the hindbrain was selected. All images were acquired using the 20X objective. The depth of acquisition was determined in each case individually; the most dorsal portion of the head containing macrophages was selected as the first plane and 10 z-slices, 2 µm apart were acquired, with an interval of 300 seconds per acquisition and a total of 100 imaging cycles. Following 17 hours 41 minutes of acquisition, maximum intensity projection of the z-slices was performed using the Zen Black software. For the purpose of analyses, macrophages within the brain were counted at each timepoint from the acquired images.

For the *rraga* injury timelapse experiments, animals at 4 dpf were anesthetized and the tip of the tail fin was cut using a new scalpel ensuring approximately same size of incision in all animals. Immediately after injury, animals were mounted in 1.5% agarose with anesthetic, and timelapse imaging was performed for 8 hours post injury. All imaging was done using a 10X objective and 10 z-slices were selected, each slice 1 µm apart, at an interval of 300 seconds per acquisition. The number of macrophages at the site of injury (regenerated fin tissue+blastema) was counted from maximum intensity projection of the z-slices. For the triple mutant injury experiment, tail tip injury was performed in wildtype animals and triple mutants and the animals were allowed to recover in embryo water. 10 hours post injury, all larvae were mounted in 1.5% agarose with anesthetic and the number of macrophages at the injury site was counted.

### Dissociation of larvae, macrophage sorting, and RNA extraction

*rraga*^*+/-*^; *mpeg:GFP* animals were intercrossed, embryos collected, treated with PTU before 24 hpf, and washed daily with embryo water containing PTU until 6 dpf. At 4 and 5 dpf, animals were anesthetized, mounted in a drop of embryo water on a slide containing coverslip bridges and examined under the confocal microscope for the presence of the GFP transgene. Animals were scored as mutants or wildtype siblings based on the morphology of macrophages (amoeboid corresponding to *rraga* mutants) and separated into dishes. At 6 dpf, larvae were anesthetized using tricaine and left at 4 ºC for 30 minutes to euthanize the animals. Larvae were washed once with cold HBSS (ThermoFisher, 14025092) and each animal was cut into at least three pieces with a fresh scalpel on ice. Slices of 90 animals of each genotype were divided into three 15 ml conical tubes (3×30 for each genotype), spun down at 800 X g for 5 minutes at 4 ºC. HBSS was removed and 2 ml of freshly made 1 mg/ml collagenase (Worthington Biochemical, LS004194) in HBSS was added to each tube. Collagenase digestion was carried out with gentle agitation in an incubator set at 32 ºC for 30 minutes; larval tissue was pipetted up and down 10X using a P1000 after 15 minutes of incubation. Following a spin at 800 X g for 5 minutes at 4 ºC, 5 ml of 0.25% (5X) cold Trypsin-EDTA (Santa Cruz Biotechnology, sc-363354, 1:1 of 0.5% or 10X in HBSS) was added to each tube. Trypsin digestion was carried out for 20 minutes with gentle agitation at room temperature; larval tissue was pipetted up and down 10X using a P1000 after 10 minutes of Trypsin digestion. Trypsin digestion was stopped by adding FBS at 5% (250 µl FBS in 5 ml Trypsin-EDTA). Tubes were spun at 800 X g for 5 minutes at 4 ºC, trypsin was removed, and the pellet was resuspended in 2 ml of cold FACSmax dissociation solution (Genlantis, T200100). Following 10 minutes of incubation in FACSmax with gentle agitation at room temperature, tubes were spun at 800 X g for 5 minutes at 4 ºC, FACSmax was removed, and the pellet was resuspended in 2 ml of cold HBSS containing 5% FBS. Following another spin, the pellet was resuspended in 250 µl of PBS containing 2% FBS and 1 mM EDTA. The pellet in PBS was strained through a 40-micron cell strainer, the cell suspension was transferred to a 5 ml FACS tube to proceed with sorting. All cell suspensions were kept on ice during the entire duration of the sort. TO-PRO-3 (AAT Bioquest 17572) was used to distinguish the live GFP^+^ cells and added at 1:1000 concentration (in PBS) 20 minutes before each tube was sorted. GFP^+^ cells (at least 10,000 cells) from *rraga* mutants or wildtype siblings were sorted using BD FACSAria II directly into lysis buffer (RLT+beta-mercaptoethanol) from Qiagen RNeasy Micro Kit (74004). RNA was extracted immediately after sorting using Qiagen RNeasy Micro Kit (74004) and the eluate containing RNA was snap frozen and stored at -80 ºC.

### Whole larvae sample preparation for RNA-Sequencing

Wildtype *mpeg:GFP* or *tfeb; tfe3a; tfe3b* triple mutants or *rraga*^*+/-*^; *mpeg:GFP* heterozygotes were intercrossed to obtain wildtype, triple mutant, and *rraga* mutant larvae respectively. At 4 dpf, approximately 100 each of wildtype or triple mutant larvae were injected with Zymosan Texas Red. After approximately one hour post injection, the uptake of Zymosan Texas Red particles by peripheral macrophages was visualized by temporarily mounting anesthetized larvae on slides with coverslip bridges and examining the animals using a confocal microscope. Animals with substantial Texas Red labeling in peripheral blood vessels, as well as amoeboid morphology of macrophages, were selected and allowed to recover in embryo water. Uninjected wildtype or triple mutants were not screened on the confocal. Approximately 2 hours post injection, 20 larvae of each genotype (wildtype or triple mutant) or treatment (uninjected or Zymosan A Texas Red injected) were pooled into snap cap tubes in triplicates (total of 60 larvae), the excess was water removed, and larvae were frozen on dry ice. *rraga* mutants were screened based on amoeboid morphology of macrophages at 4 dpf as described above and snap frozen on dry ice. All the larvae were homogenized using a FastPrep-24 Classic bead beating grinder and lysis system (MP Bio), RNA was extracted using Takara NucleoSpin RNA Plus (740984.50) kit and the eluate containing RNA was snap frozen and stored at -80 ºC.

### Library Preparation and Quality Control

The subsequent steps of RNA-Sequencing were performed by Novogene Corporation Inc. Quality control for all RNA samples was performed using Agilent 2100 Bioanalyzer system; all samples had a sample integrity value of RIN>9.0. For the macrophage RNA-Seq experiment, five biological replicates were initially processed; following RNA quality control, four biological replicates were advanced for library preparation and sequencing. For whole larvae sequencing, three biological replicates were used for each treatment and/or genotype. cDNA library preparation was done using Takara SMART-Seq v4 Ultra Low Input RNA Kit for the macrophage RNA samples and NEBNext Ultra II RNA Library Prep Kit for the whole larvae RNA samples. Sequencing was performed using Illumina NovaSeq platform (PE150). Data quality control was performed to confirm that the error rate was 0.02-0.03% for all samples and GC content of paired reads was approximately 50%. At least 82 million reads were obtained for each of the macrophage samples and over 39 million reads were obtained from whole larvae samples.

### RNA-Sequencing analysis

After reads containing adapters and low-quality reads were removed, the percentage of clean reads was over 94% from macrophage samples and at least 97% for samples from whole larvae. These reads were mapped using HISAT2 v2.0.5 to Ensembl GRCz11 (Genome Reference Consortium Zebrafish Build 11) and quantification of reads was done using featureCounts v1.5.0 (*72*). Over 90% of the clean reads were mapped to the genome and close to 80% of these reads mapped to exonic regions in the genome. Correlation analysis was performed to confirm that biological replicates had a Pearson correlation coefficient of >0.92 and Principal Component Analysis was performed to ensure that biological replicates clustered together. Differential expression analysis was done using DESeq2 R package (*73*), and the resulting P values were adjusted using the Benjamini and Hochberg’s approach for controlling false discovery rate. A padj cut off of <0.05 and log_2_fold change >+1 (for differentially upregulated genes) or <-1 (for differentially downregulated genes) was used for all samples with the exception of the heatmap in Figure 7. ClusterProfiler (*74*) and ShinyGO (*75*) was used for Gene Ontology (GO) and Kyoto Encyclopedia of Genes and Genomes (KEGG) enrichment analyses. Fold Enrichment in GO enrichment graphs is defined as the percentage of genes in your list belonging to a pathway, divided by the corresponding percentage in the background. Qiagen Ingenuity Pathway Analysis software was used to determine top canonical pathways enriched in differentially expressed genes.

### EdU labeling

*rraga*^*+/-*^; *mpeg:GFP* animals were intercrossed and dechorionated at approximately 50 hpf. Larvae were anesthetized and 1 nanoliter of 5 mM EdU from ThemoFisher (C10086, 10 mM EdU stock in DMSO was diluted 1:1 in 1X PBS before injections) was injected in the duct of Cuvier at 60 hpf. The EdU was chased overnight for 12 hours and animals were fixed at 72 hpf with 4% paraformaldehyde for 2 hours at room temperature. Fixed larvae were permeabilized using Proteinase K (ThermoFisher, 25530049) at a dilution of 1:1000 for 30 minutes and post-fixed using 4% paraformaldehyde for 20 minutes at room temperature. Larvae were incubated in blocking solution (1% DMSO, 1% donkey serum, 1% BSA, 0.7% Triton X-100 in 1X PBS) for 2 hours at room temperature and incubated in blocking solution with anti-GFP antibody (1:500, Abcam, ab6658) overnight at 4 ºC. Following 6 10-minute washes in 1X PBS with 0.8% Triton X-100, larvae were incubated in block solution with 1:500 donkey anti-goat AlexaFluor 488 or 594 overnight at 4 ºC. After 6 10-minute washes in 1X PBS with 0.8% Triton X-100, Click-iT reaction was performed according to manufacturer’s instructions (ThermoFisher, C10086) and animals were incubated in development solution for 1 hour in the dark at room temperature. Following washes with 1X PBS with 0.8% Triton X-100 overnight, animals were mounted in 80% glycerol and imaged. Quantification and genotyping were performed post-imaging.

### Yolk injection of dyes or debris to assay macrophage uptake

Injections of DQ Red-BSA (ThermoFisher D12051, 1 mg/ml in 1X PBS), Magic Red Cathepsin (Immunochemistry Technologies 937, vial 6133 suspended in 50 µl DMSO, 1:1 dilution in 1X PBS before use), *E. coli* Texas Red (ThermoFisher E2863, 20 mg/ml in 1X PBS), Zymosan (Sigma Aldrich Z4250, 10 mg/ml in 1X PBS, boiled to solubilize), and Zymosan Texas Red (ThermoFisher Z2843, 20 mg/ml in 1X PBS) were all performed at 4 dpf. 1 nl of each of the above suspensions was injected in the yolk of anesthetized larvae (*76*) of the required genotype. The injected dyes or debris particles migrate to the peripheral blood vessels of the larvae within approximately 1-1.5 hours post injection, where macrophages are easily visualized. For microscopy experiments, larvae were anesthetized, mounted in 1.5% agarose, and imaged using Zeiss LSM Confocal microscope. For RNA-Sequencing experiments, larvae injected with Zymosan Texas Red were mounted in a drop of embryo water on a slide containing coverslip bridges and returned to embryo water to recover after the uptake of Texas Red particles by macrophages was confirmed.

### Transgene construct and injection

Full length zebrafish *tfe3b* (accession number: NM_001045066.1) and *LAMP1b* (accession number: NM_001326532.1) were cloned from wild type cDNA and *LAMP2, rab5*, and *rab7* were amplified and cloned from respective plasmids (*47, 77*). All the cloning primers are listed in Table S1. Amplified PCR fragments of respective genes were cloned into pCR8 vector (for *tfe3b*) or pCR8 vector with mCherry (for *LAMP1b, LAMP2, rab5*, and *rab7*) and verified by Sanger sequencing. Macrophage-specific expression vector was assembled using multisite Gateway method using the *mpeg* promoter and a destination vector containing Tol2-transposon sites for genomic integration and *Tg(cmlc2:mCherry)* or *Tg(cmlc2:GFP)* for selection (*78*). The plasmid expressing the transgene was co-injected at 12-25 pg along with 50-100 pg of Tol2 transposase mRNA at 1-cell stage (*78*). For *tfe3b*, injected embryos were selected for further analysis based on strong expression of mCherry in the heart by the cardiac reporter *Tg(cmlc2:mCherry)*. The injected animals were raised, outcrossed to TL or *Tg(mpeg:GFP)* lines, and presence of stable expression of mCherry in the heart was confirmed prior to experiments and/or imaging. *Tg(cmlc2:GFP)* was used as the selection marker for *lamp1b-mCherry, lamp2-mCherry, mCherry-rab5*, and *mCherry-rab7* constructs. The Rab5 and Rab7 transgene constructs were crossed to *rraga*^*+/-*^; *mpeg:GFP* lines and microscopy experiments were performed on F1 generation. LAMP1B and LAMP2 experiments were performed on *rraga*^*+/-*^; *mpeg:GFP* intercross animals after injection and normalization for *cmlc2:GFP* expression (F0 generation).

### Generation of *tfe3a, tfe3b*, and *flcn* mutants with CRISPR-Cas9

sgRNAs targeting *tfe3a, tfe3b*, and *flcn* were designed using CHOPCHOP (https://chopchop.cbu.uib.no/) (*79*). Oligonucleotides containing T7 binding site and the CRISPR sequence were annealed to tracrRNA template. Assembled oligonucleotides were transcribed using HiScribe T7 Quick (NEB, E2050S) kit. Following DNase treatment, the RNA was purified using mirVana miRNA isolation kit (Invitrogen, AM1561). An aliquot of the sgRNA eluate was run on agarose gel and quantified using NanoDrop 8000. CRISPR injections were performed at 1-cell stage. Injection mix consisted of 300 ng/µl of Cas9 protein (Macrolab, Berkeley, http://qb3.berkeley.edu/macrolab/cas9-nls-purified-protein/) and 300 ng/µl of the sgRNA in Tris-HCl (pH 7.5). A small amount of phenol red was added to the mix to help with visualization during injection. Injected fish were raised to adulthood.

The F1 progeny of F0-injected fish were screened for out-of-frame insertions or deletions. Details of guideRNAs used, lesions, and restriction enzyme assays for genotyping mutants are described in Table S1. The *tfe3a* allele, *st124*, is a 2 bp deletion in exon 2. We used two mutant alleles of *tfe3b: st125*, a 25 bp deletion in exon 2, and *st126*, which is linked to *tfeb* (*st120)*, and is a 2 bp deletion in exon 2. The *flcn* mutation has two different lesions generated by co-injecting two guide RNAs: *st127* has a 7 (-8+1) bp change in exon 2 (which disrupts the open reading frame), followed by a 4 bp deletion in exon 7, approximately 10,000 kb downstream.

### Neutral red assay

Neutral red was dissolved in distilled water to give a 2.5 mg/ml stock solution, which is stable at room temperature for several months. Staining using neutral red was done by treating larvae at 4 dpf with 5 µg/ml solution of neutral red in embryo water containing PTU for 3 hours (*44*). Animals were rinsed at least twice after neutral red staining and washed overnight in embryo water with PTU at 28.5 ºC. Approximately 24 hours post-neutral red treatment, larvae were anesthetized and mounted in 1.5% low melting point agarose. The number of neutral red^+^ microglia were counted, and images were acquired immediately after counting using Zeiss AxioCam HRc camera with the AxioVision software. Between 100-150 embryos were mounted and imaged for each double, triple, and quadruple mutant experiment. After imaging, all animals were genotyped by PCR and statistical analysis was performed.

### LysoTracker Red and LysoSensor Green assays

LysoTracker Red DND-99 (ThermoFisher, L7528) was diluted 1:100 in embryo water+PTU and LysoSensor Green DND-189 (TheroFisher, L7535) was diluted 1:1000 in embryo water+PTU. Larvae at 4 dpf were incubated in the LysoTracker Red or LysoSensor Green solution for 30-45 minutes. After staining, larvae were washed in embryo water containing PTU for 30 minutes, with at least 3 washes approximately 8-10 minutes apart. Following anesthesia and mounting in agarose, images were acquired using Zeiss LSM confocal microscope. All genotyping was done by PCR post-imaging and intensity of LysoSensor Green or area of LysoTracker Red punctae in the midbrain, where most microglia are present, was calculated using ImageJ for at least three animals per genotype.

### in situ hybridization

*apoe* antisense probe was generated from a pCRII clone carrying 505 bp of *apoeb* gene (*40*). in situ hybridization was performed using standard methods (*81*). Briefly, embryos at 4 dpf were fixed overnight in 4% paraformaldehyde, dehydrated at least overnight in 100% methanol, rehydrated in PBS-Triton X-100 and PBS, permeabilized using proteinase K (20 mg/ml) at a dilution of 1:1000 for 1 hour, and incubated overnight with antisense riboprobes at 65 ºC. Following washes in 2X and 0.2X SSC, and incubation in MAB block containing 10% normal sheep serum, animals were incubated overnight at 4 ºC in MAB block containing 1:1000 dilution of anti-digoxigenin antibody conjugated to alkaline phosphatase. The following day, after at least 6 20-minute washes in MAB-Triton X-100 buffer, animals were incubated in development solution containing NBT (Roche, 11383213001) and BCIP (Roche, 11383221001). Development was stopped at the same time for all samples in a single experiment by evaluating the strength of the *apoe* signal in control animals. Animals were washed twice in PBS-Triton X-100, left in 100% ethanol overnight for destaining, and rehydrated in PBSTx. Following genotyping of finclips, representative animals were mounted in 100% glycerol and images were captured using Zeiss AxioCam HRc camera with the AxioVision software.

## Acknowledgments

We thank Tuky Reyes and Chenelle Hill for maintaining the fish facility. We are grateful to Daniel Lysko, Mariapaola Sidoli, Ellen Bouchard, and John Vaughen for their thoughtful comments on the manuscript. Templates for the constructs generated in this manuscript were generously donated by colleagues: Rab5 and Rab7 plasmids from Brian Link at the Medical College of Wisconsin, LAMP2 plasmid from Michel Bagnat at Duke University, and mCherry plasmid from Garrett Kingman at Stanford University.

## Funding

Postdoctoral Fellowship (18POST33990334), American Heart Association (HI)

Developmental Project Award, Stanford Alzheimer’s Disease Research Center and National Institute on Aging (HI)

Postdoctoral Fellowship (A2021011F), BrightFocus Foundation (HI)

A^∗^STAR Singapore (KS)

National Institutes of Health grant R35NS111584 (WST, AMM)

National Multiple Sclerosis Society grant RG-1707-28694 (WST, AMM)

## Author contributions

Conceptualization: HI

Methodology: HI, WST I

Investigation: HI, KS, AMM

Visualization: HI

Supervision: WST

Writing – original draft: HI

Writing – review and editing: HI, WST, AMM

## Data and material availability

All data are available in the main text or the supplementary materials.

