## Supplemental figures for "Repression of lysosomal transcription factors Tfeb and Tfe3 is essential for the migration and function of microglia"

Fig. S1

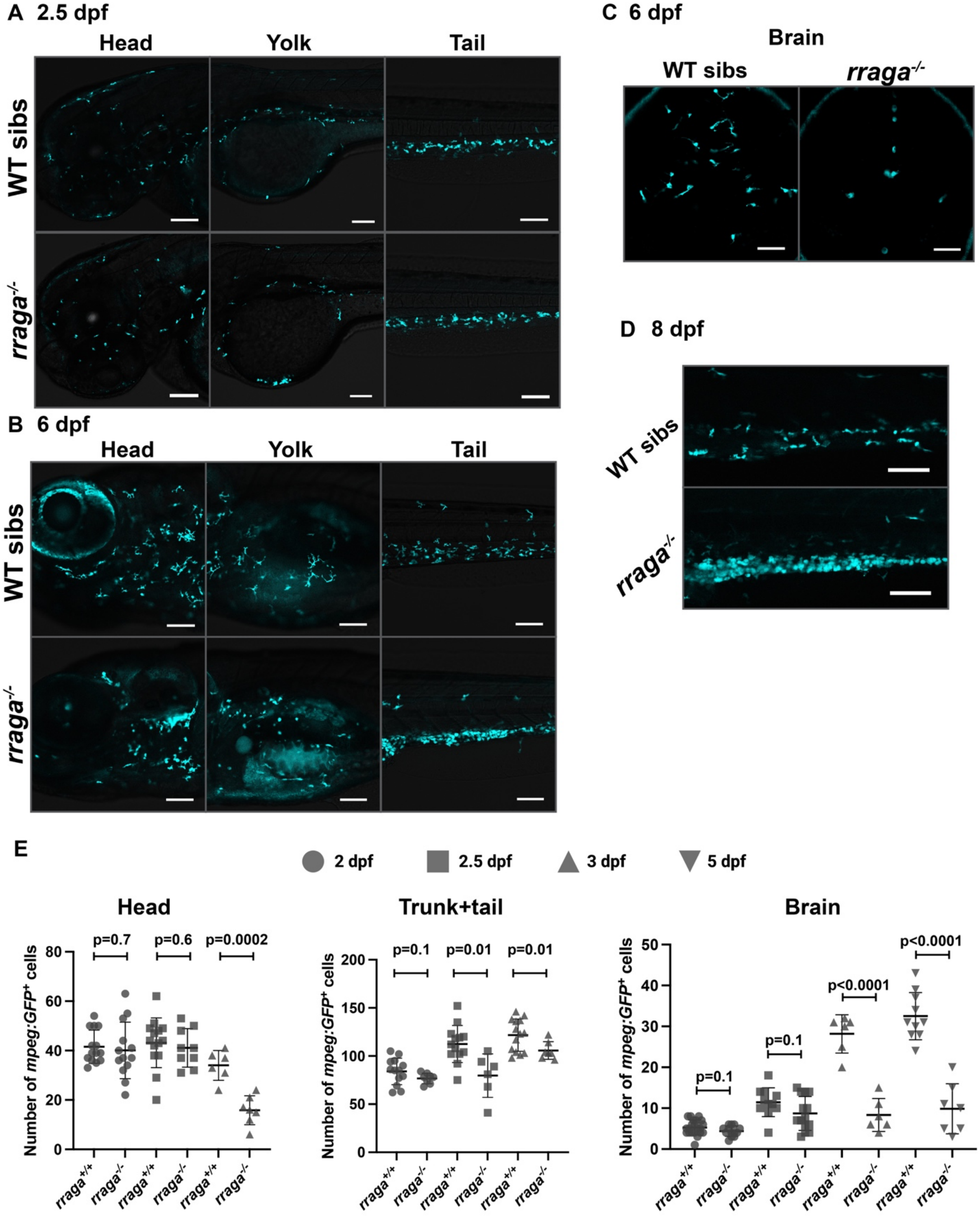


**Fig. S1. Abnormal peripheral macrophages persist in *rraga* mutants.** Images of *mpeg:GFP*-expressing macrophages and microglia in *rraga* mutants and their wildtype siblings at (A) 2.5 dpf, (B, C) 6 dpf, and (D) 8 dpf. (A, B, D) show lateral views of the animal with anterior on top. Scale bars, 100 µm. (D) Dorsal views of the brain. Scale bars, 50 µm. (E) Quantification of GFP$^+$ cells in images from Figures 1I-L and S1A. All graphs show mean with standard deviation; significance was determined using unpaired t-test.

Fig. S2

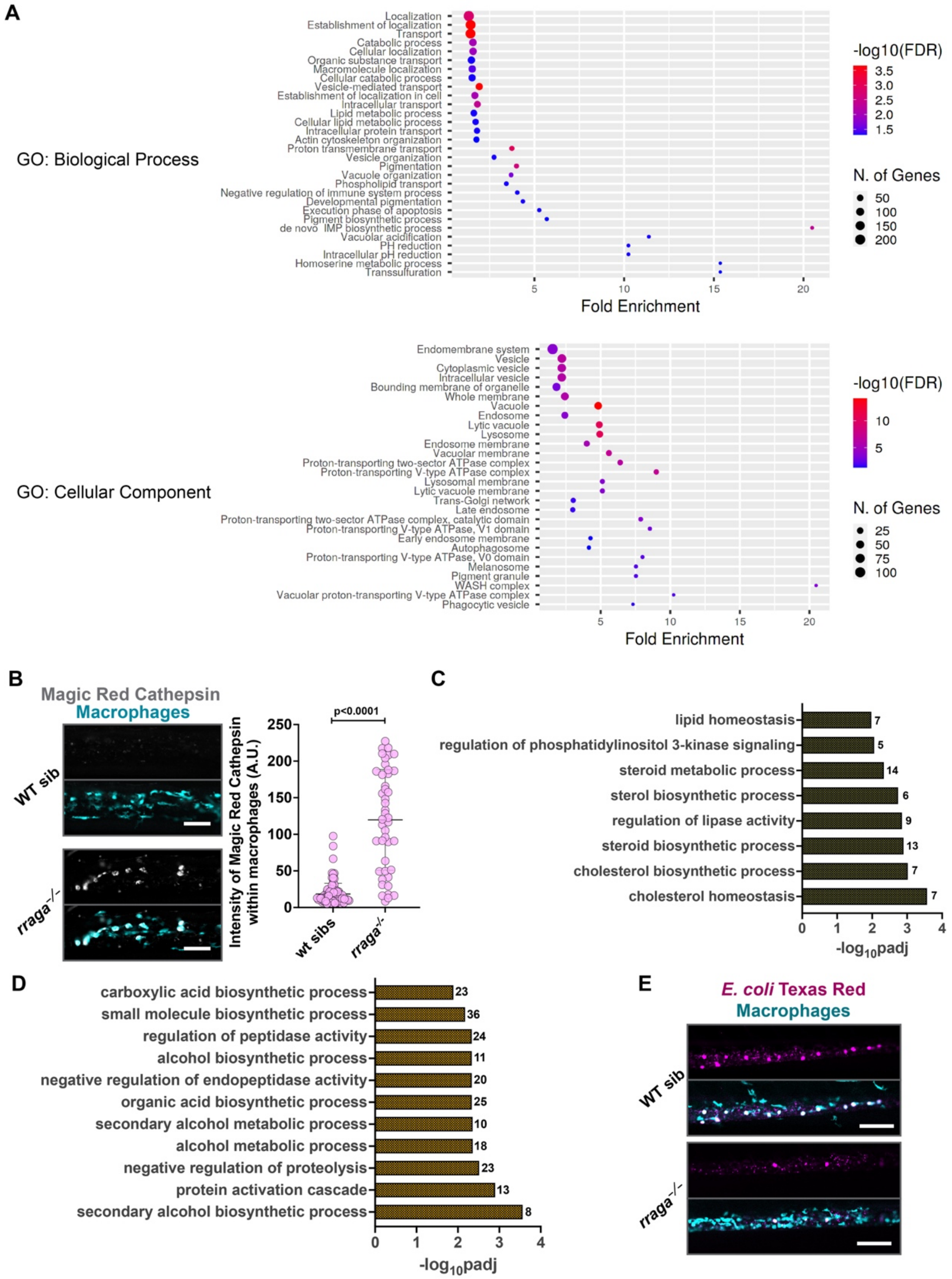


**Fig. S2. Functional validation of RNA-Seq and *rraga* mutants.** (A) Graphs showing GO term enrichment analysis (Biological Process and Cellular Component) of genes upregulated in macrophages from *rraga* mutants. (B) Magic Red Cathepsin labeling and quantification to detect proteolytic activity in macrophages. GO terms corresponding to (C) lipid metabolism and (D) other metabolic pathways were also significantly downregulated in macrophages from *rraga* mutants. (E) Low magnification view of the tail showing the extent of uptake of *E. coli* Texas Red particles by macrophages.

Fig. S3

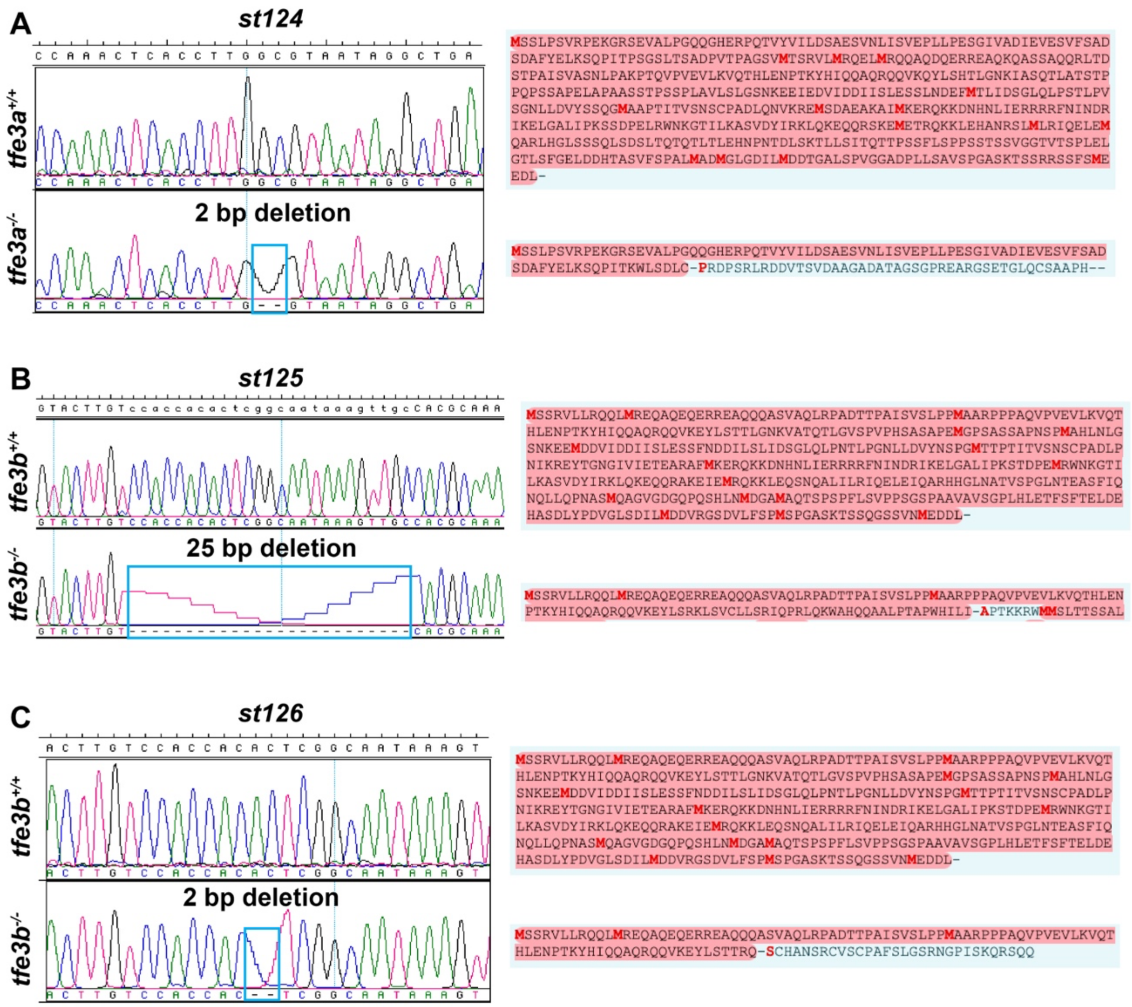


**Fig. S3. *tfe3a* and *tfe3b* deletions.** Chromatogram of wildtype and mutant sequences and predicted sequences of corresponding proteins in (A) *tfe3a, st124,* 2bp deletion in exon 2; (B) *tfe3b, st125,* 25 bp deletion in exon 2; and (C) *tfe3b, st126,* 2 bp deletion in exon 2 (on the same chromosome as *tfeb, st120*).

Fig. S4

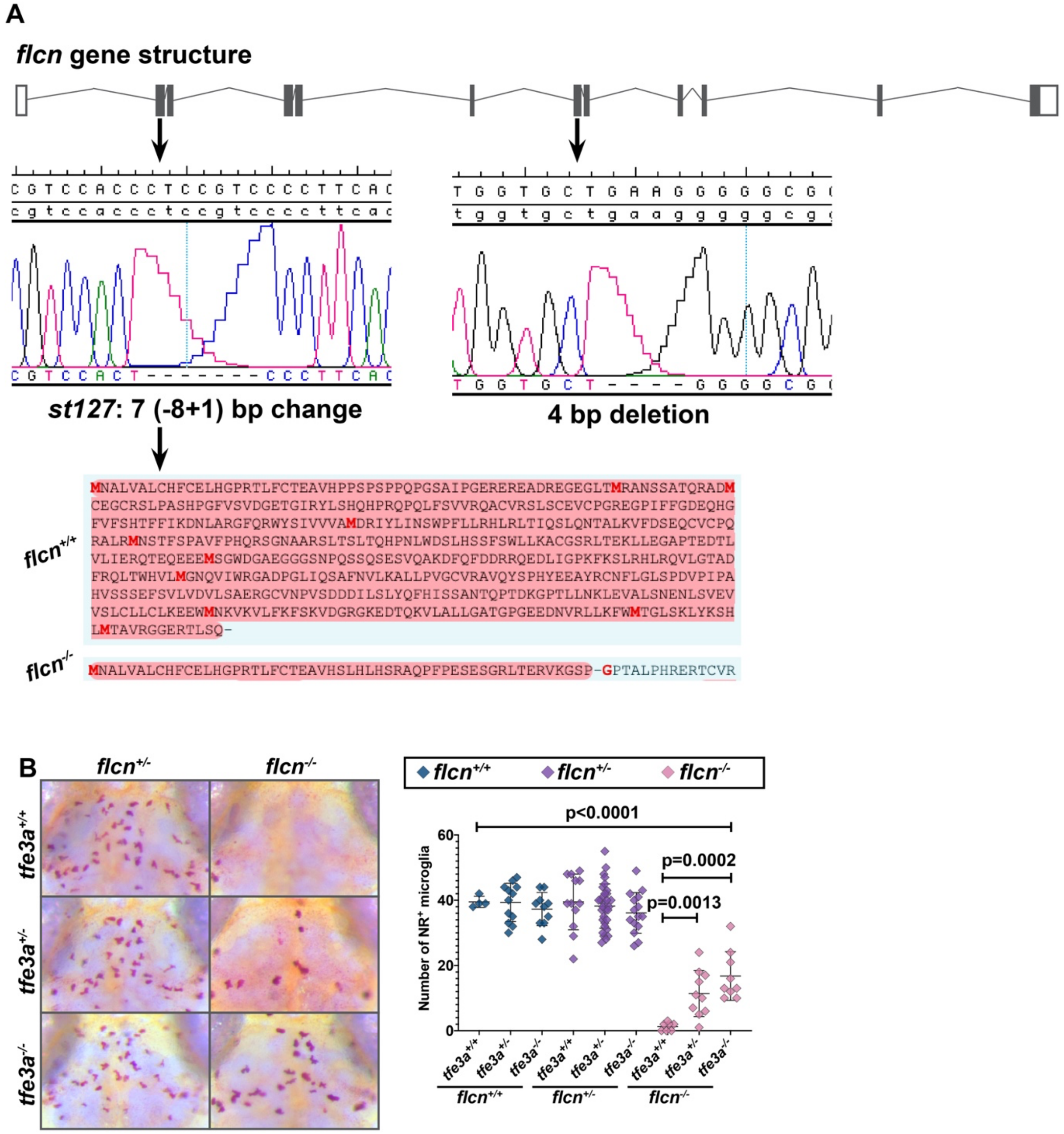


**Fig. S4. *flcn* mutation and mutant phenotypes.** (A) *flcn* gene structure and chromatograms showing sequence changes in *st127.* Gene structure diagram not to scale. (B) NR assay and quantification of microglia in progeny of *flcn*$^{+/-}$; *tfe3a*$^{+/-}$ intercross.

**Fig. S5**

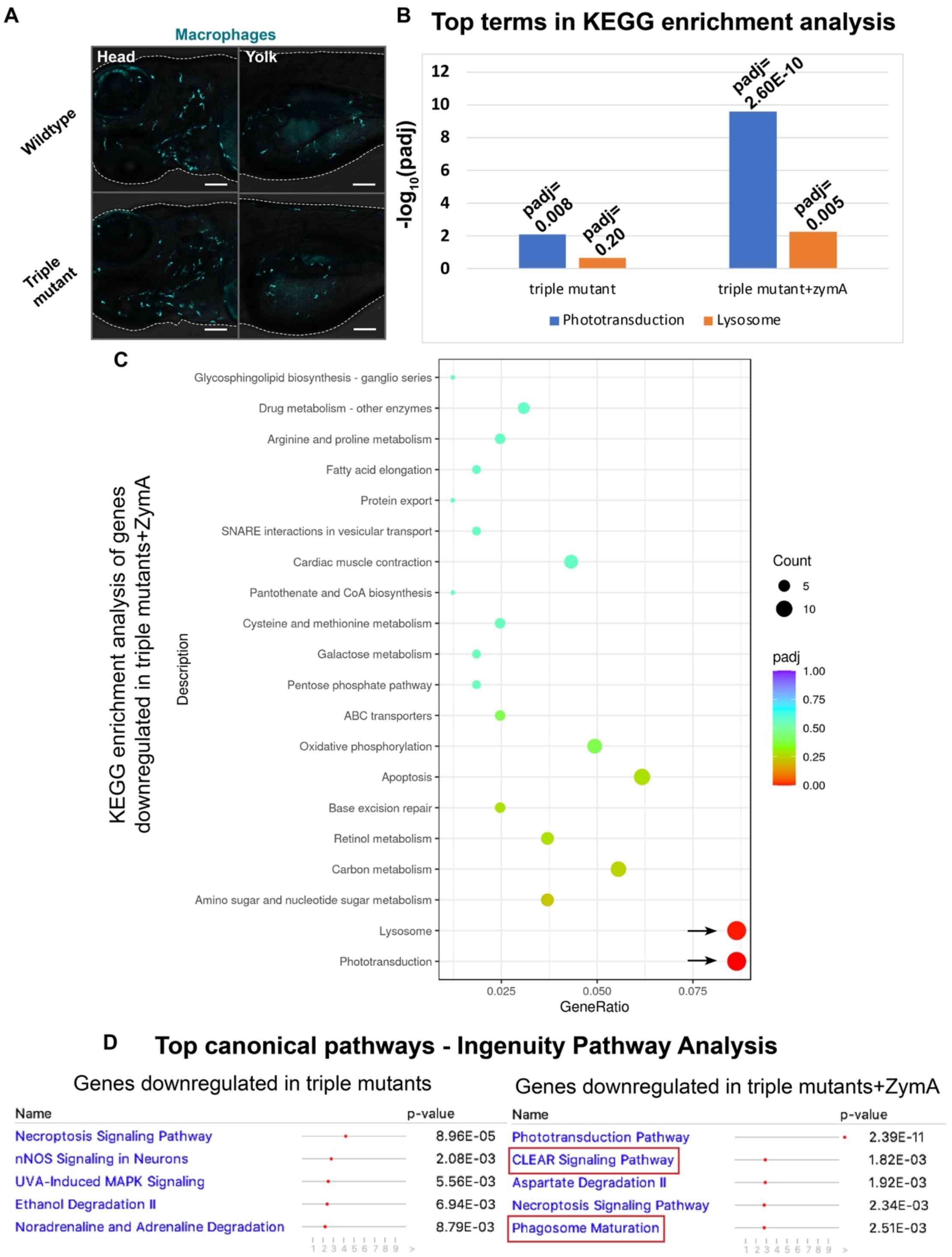


**Fig. S5. Tfeb and Tfe3 activate lysosomal pathways only under conditions of stress.** (A) Macrophages visualized using the *mpeg:GFP* transgene in wildtype animals and triple mutants. (B) Graph depicting the padj values of the top terms in KEGG enrichment analysis of genes significantly downregulated in triple mutants injected with ZymA and uninjected triple mutants. No other KEGG pathway shows a significant difference (padj<0.05). (C) Dot plot of KEGG pathways analysis of genes significantly downregulated in triple mutants injected with ZymA. (D) Top canonical pathways following Ingenuity Pathway Analysis of genes significantly downregulated in triple mutants or triple mutants injected with ZymA.

**Table S1. Details of CRISPR mutants described in this manuscript.**

**Table S2. GO term enrichment analysis on genes upregulated in macrophages isolated from *rraga* mutants**

**Table S3. GO term enrichment analysis on genes downregulated in macrophages isolated from *rraga* mutants**

**Table S4. GO enrichment analysis on differentially expressed genes in triple mutants versus wildtype**
